# 6′,6′-Difluoro-aristeromycin is a potent inhibitor of MERS-coronavirus replication

**DOI:** 10.1101/2021.05.20.445077

**Authors:** Natacha S. Ogando, Jessika C. Zevenhoven-Dobbe, Dnyandev B. Jarhad, Sushil Kumar Tripathi, Hyuk Woo Lee, Lak Shin Jeong, Eric J. Snijder, Clara C. Posthuma

## Abstract

The severe acute respiratory syndrome coronavirus 2 (SARS-CoV-2) pandemic has highlighted the lack of treatments to combat infections with human or (potentially) zoonotic CoVs. Thus, it is critical to develop and evaluate antiviral compounds that either directly target CoV functions or modulate host functions involved in viral replication. Here, we demonstrate that low-micromolar concentrations of 6′,6′-difluoro-aristeromycin (DFA), an adenosine nucleoside analogue, strongly inhibit the replication of Middle East respiratory syndrome coronavirus (MERS-CoV) in a cell-based infection assay. DFA was designed to target S-adenosylhomocysteine (SAH) hydrolase and, consequently, may affect intracellular levels of the methyl donor S-adenosylmethionine, which is used by two CoV methyltransferases involved in the capping of the 5’ end of the viral mRNAs. Passaging of wild-type MERS-CoV in the presence of DFA selected a virus population with a ∼100-fold decreased DFA sensitivity, which carried various amino acid substitutions in viral nonstructural proteins (nsps). Specifically, mutations were present in the RNA polymerase subunit (nsp12) and in nsp13, the helicase subunit containing a nucleoside triphosphate hydrolase activity that has been implicated in CoV capping. We hypothesize that DFA directly or indirectly affects viral cap methylation, either by inhibiting the viral enzymes involved or by binding to SAH hydrolase. We also evaluated the antiviral activity of DFA against other betacoronaviruses, but found it to have limited impact on their replication, while being quite cytotoxic to the Calu-3 cells used for this comparison. Nevertheless, our results justify the further characterization of DFA derivatives as an inhibitor of MERS-CoV replication.

**Importance:** Currently, there is a lack of antiviral drugs with proven efficacy against human CoV infections including the MERS-CoV that is endemic in the Middle East, the pandemic SARS-CoV-2 and potential future zoonotic CoV. This highlights the importance to investigate new drug targets and identify compounds that can be used to inhibit CoV replication. In this study, we characterize the inhibitory effect of DFA on MERS-CoV replication by phenotypic studies, time-of-addition studies, and the generation and genotyping of a DFA-resistant virus population. Our results revealed that DFA needs further improvement to reduce its cytotoxic side-effects and potentially enhance its broad-spectrum activity. Despite this observation, we think that DFA can be used to understand the function and metabolic interactions of the CoV RNA-synthesizing machinery, or as a starting point for the design of new compounds of the same class.

## Introduction

Previously, the emergence of severe acute respiratory syndrome coronavirus (SARS-CoV; in 2003 in China) and Middle East respiratory syndrome coronavirus (MERS-CoV; in 2012 in Saudi Arabia) highlighted the potential pandemic threat posed by this type of zoonotic pathogens and the need to develop rapid response options to contain them (1-4). Due to the severity of the diseases caused by SARS-CoV and MERS-CoV, and their potential for zoonotic transmission and global spread, both these agents received a priority status from the World Health Organization and other government agencies for the development of prophylactic and therapeutic treatment strategies (5, 6). The current SARS-CoV-2 pandemic (7, 8) and its burden on public health worldwide further emphasize the critical nature of the quest for anti-CoV drugs with high clinical efficacy. Many drug classes currently are under evaluation as inhibitors of CoV replication, including compounds directly targeting viral functions, like viral proteases and the RNA polymerase, and host factor-targeting inhibitors (reviewed in (9-12)).

Coronaviruses are positive-stranded RNA (+RNA) viruses with a single genomic RNA of approximately 30 kb that is replicated in the cytoplasm of infected cells. Following entry, the 5’-capped viral genome is recognized and translated by host ribosomes to yield the replicase polyproteins pp1a and pp1ab (13). Subsequently, these large precursors are processed into 16 individual nonstructural proteins (nsp 1 to 16), which are released following polyprotein cleavage by two or three internal proteases. Together, the nsps form a multi-enzyme complex that ensures the replication of the viral genome and the transcription of a set of subgenomic mRNAs (reviewed in (14, 15)). The enzymatic core of this complex is formed by the nsp12 RNA-dependent RNA polymerase (RdRp) that synthesizes RNA with the help of the auxiliary factors nsp7 and nsp8 (16, 17), the nsp13 helicase that unwinds RNA duplexes (18-20), and several other RNA-processing enzymes residing in nsp12-nsp16 (reviewed in (15, 21, 22)). These also include a 3’-to-5’ exoribonuclease (nsp14-ExoN) that is thought to increase replication fidelity by correcting mismatches sustained during RNA synthesis (reviewed in (23-26)). The viral structural and accessory proteins, encoded by smaller open reading frames located in the 3’-proximal part of the genome, are expressed from a set of 5’-capped and 3’-polyadenylated subgenomic mRNAs (reviewed in (15, 22)). Apart from ensuring mRNA recognition during formation of the ribosomal preinitiation complex, the 5’-terminal cap structure protects the viral mRNAs from degradation by cellular ribonucleases and prevents detection by the host’s intracellular pathogen recognition receptors, which would trigger innate immune responses (reviewed in (27)).

The CoV capping mechanism is thought to consist of four sequential reactions: (i) an RNA triphosphatase activity residing in nsp13 removes the γ-phosphate group from the 5′-triphosphorylated RNA (28, 29); (ii) a guanosine monophosphate (GMP) is transferred to the 5’-diphosphate terminus by a yet to be confirmed guanylyltransferase (GTase)(30), which was recently proposed to reside in the N-terminal nucleotidyl transferase (NiRAN) domain of nsp12 (31); (iii) the nsp14 methyltransferase (MTase) methylates the cap’s 5’-terminal guanine at the N7-position, producing the so-called cap-0 structure, ^7me^GpppN (32); (iv) finally, a cap-1 structure is formed when nsp16, in complex with its nsp10 co-factor, methylates the ribose 2’-O-position of the first transcribed nucleotide of each viral RNA, converting ^7me^GpppN into ^7me^GpppN_2’me_ (33). Given the central position of the RNA-synthesizing and capping machinery in the CoV replication cycle, each single component constitutes a potential target for direct-acting antiviral drug development.

As in cellular methylation reactions, S-adenosyl-L-methionine (SAM) is the most common methyl donor used by viral MTases, such as those present in CoV nsp14 and nsp16 (34, 35). Thus, the identification of compounds that can interfere with viral mRNA capping, by either directly targeting viral MTases or indirectly affecting the concentrations of essential cellular metabolites, constitutes a viable strategy to develop broad-spectrum CoV inhibitors. S-Adenosyl-homocysteine (SAH) is released upon the transfer of the methyl group of SAM to a nucleic acid substrate by a SAM-dependent MTase. Consequently, accumulation of SAH can interfere with SAM-dependent MTase function due to product inhibition (36). Inhibitors targeting S-adenosyl-homocysteine (SAH) hydrolase have been reported as potential broad-spectrum antiviral drugs in different studies (37-39). This hydrolase catalyzes the reversible conversion of SAH into adenosine and L-homocysteine, which both are then further metabolized for use in different cellular pathways (40, 41).

Recently, using cell-based assays for MERS-CoV, SARS-CoV, chikungunya and Zika virus replication, we described the inhibitory potential of a set of adenosine and selenoadenosine analogues (38). These compounds were derived from aristeromycin, a well-known carbocyclic nucleoside compound that inhibits SAH hydrolase and exhibits anti-viral, anti-cancer and anti-toxoplasma activities (reviewed in (42)). These aristeromycin derivatives are nucleoside analogues designed to directly target viral RdRp activity and/or indirectly target the methylation of viral RNA by inhibiting the host SAH hydrolase (38). From this library, we identified 6′,6′-difluoro-aristeromycin (DFA) as the aristeromycin derivative that inhibited MERS-CoV replication most efficiently in cell-based assays (38). In different cell lines, DFA inhibited MERS-CoV replication at low-micromolar concentrations and could potently reduce the progeny titers produced by MERS-CoV. Evaluation of the potential of DFA as a broad-spectrum antiviral compound revealed limited inhibition of the replication of different betacoronaviruses at non-cytotoxic concentrations. This suggests that DFA-based derivatives need to be developed to improve the antiviral activity of this compound class and reduce the cytotoxic side-effects.

## Results

### DFA inhibits MERS-CoV replication at low-micromolar concentrations in different cell lines

DFA was part of a library of more than 80 adenosine and selenoadenosine analogues that was previously evaluated for its antiviral activity against MERS-CoV, SARS-CoV and mouse hepatitis virus (MHV) using cell-based cytopathic effect (CPE) reduction assays. From this analysis, DFA was identified as the most potent inhibitor of MERS-CoV and SARS-CoV replication, with EC_50_ values (half-maximum effective concentration) of 0.2 µM and 0.5 µM, respectively. The compound was found to be more effective in reducing the progeny titers of MERS-CoV than those of SARS-CoV, yielding reductions of more than 3 log_10_ and 1 log_10_, respectively, when treating Vero cells with 1.2 µM of DFA (38). Here, we evaluated the antiviral activity of DFA against MERS-CoV in more detail using two independent cell-based assays: a CPE-reduction assay and a dose response assay, using previously described protocols (43, 44). Both assays were performed in different cell lines of human (Huh7 and MRC-5) and non-human origin (Vero).

Remdesivir (RDV) and chloroquine (CHO) were included as positive controls for inhibition of viral replication. The mean EC_50_ values in Vero cells for RDV and CHO were 0.4 µM and 25 µM, respectively, similar to what was described previously (45, 46). Using CPE reduction assays, EC_50_ values in the low-micromolar range were measured for DFA in each of the three cell lines: 0.2 µM in Vero cells, 5.2 µM in Huh7 cells, and 2.3 µM in MRC-5 cells (Fig. 1A-C). In cytotoxicity control studies, the corresponding CC_50_ values (the compound concentration resulting in 50% cytotoxicity) were calculated to be 3.6 µM in Vero, 64 µM in Huh7, and >100 µM in MRC-5 cells (Fig 1A-C). Differences between cell lines in sensitivity (cytotoxicity) to DFA treatment, as observed here, may reflect variation in SAH hydrolase expression (a target of DFA) or uptake and metabolization of the compound.

**Figure 1.**
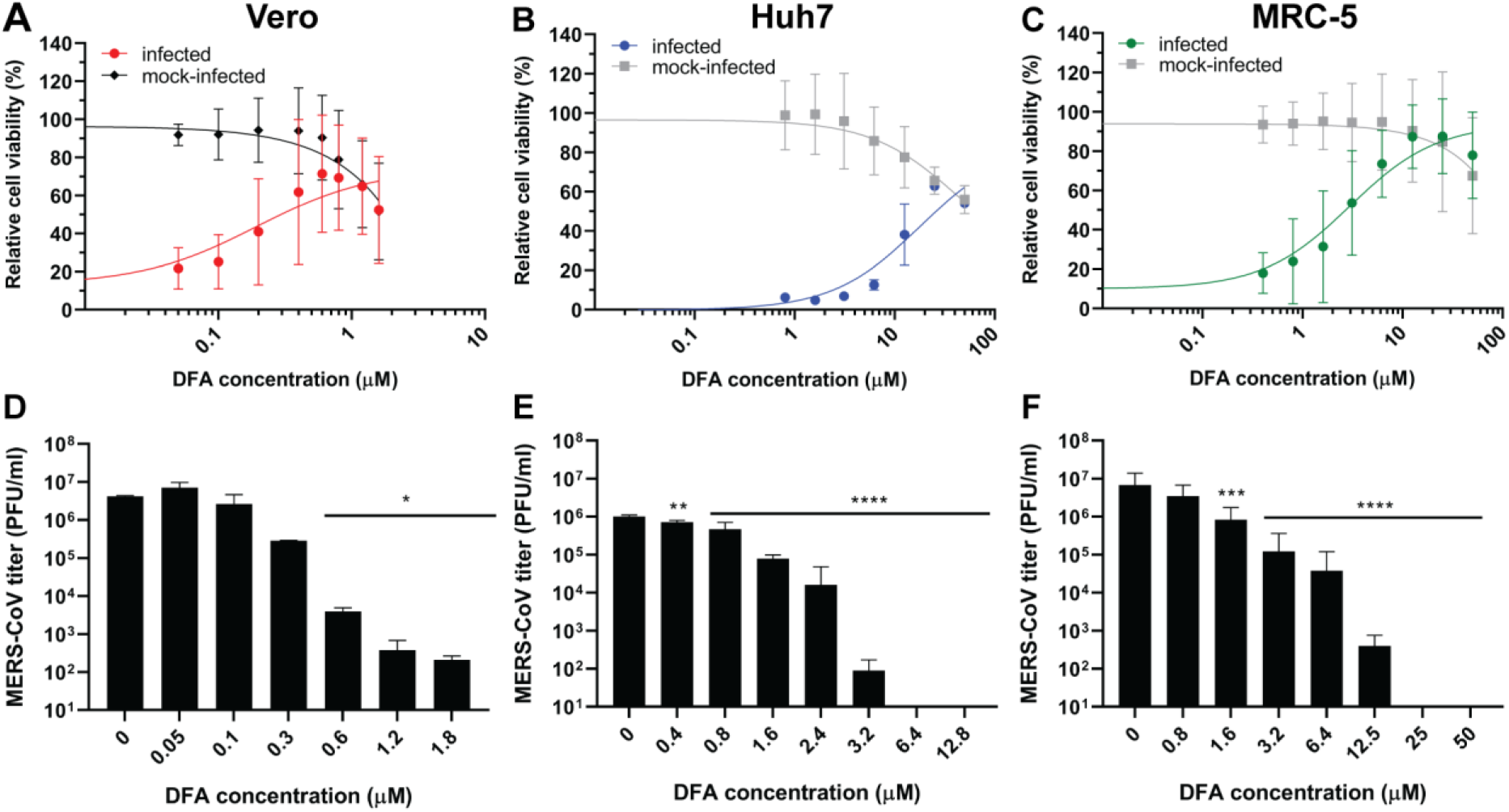
DFA inhibits MERS-CoV replication in different cell lines. Vero (A and D), Huh7 (B and E) and MRC-5 (C and F) were treated with a two-fold dilution series of DFA in the low-micromolar range and infected with MERS-CoV. Inhibitory effect was evaluated by a CPE-reduction assay (A-C) or dose response assay (D-F). For the CPE-reduction assay, cell viability was assayed using the CellTiter 96 Aqueous One Solution cell proliferation assay (MTS assay) 3 d p.i.. The graphs show the results of at least two independent experiments (mean ± sd are shown). A non-linear regression analysis was applied. In the dose response assay, cell supernatants were collected after 2 d.p.i and viral progeny was titrated by plaque assay on Vero cells. Error bars represent standard deviation. Statistical significance was determined by one-way ANOVA. *, p<0.1; **, p<0.01; ***, p<0.001; ****, p<0.0001.

In order to analyze the inhibitory effect of DFA on MERS-CoV progeny production in more detail, a multiple-cycle dose response assay was performed. A dose-dependent reduction of viral progeny was observed, with a 4 to 5 log_10_ decrease following treatment with >1.2 µM of DFA in Vero cells, >2.4 µM in Huh7, and >12.5 µM in MRC-5, respectively (Fig. 1D-F). Similar or lower EC_50_ values than in the CPE reduction assay were calculated from these studies: 0.2 µM in Vero cells, ∼0.8 µM in Huh7 cells, and 1.4 µM in MRC-5 cells. These results indicated that DFA exhibits a similar antiviral activity across multiple cell lines resulting in a consistent ∼3.5 to 4-log_10_ reduction of MERS-CoV progeny titers.

Having established the strong inhibition of MERS-CoV replication by DFA, we also tested its monophosphoramidate pro-drug (pDFA; Fig. 2A) in a CPE-reduction assay. This compound was synthesized in order to circumvent the rate-limiting first phosphorylation step that presumably restricts the efficient metabolization of nucleoside analogues like DFA following their uptake by the cell (reviewed in (47)). Unfortunately, in this case the pro-drug was less active than DFA itself, independent of the cell line used (Fig. 2B). Although the chemical and structural modifications of the prodrug decreased its cytotoxicity, the calculated EC_50_ values, 9 µM in Vero cells and 36 µM in MRC-5 cells, were more than 10 times higher than the ones measured for DFA (Fig. 1). Therefore, pDFA was not included in subsequent experiments.

**Figure 2.**
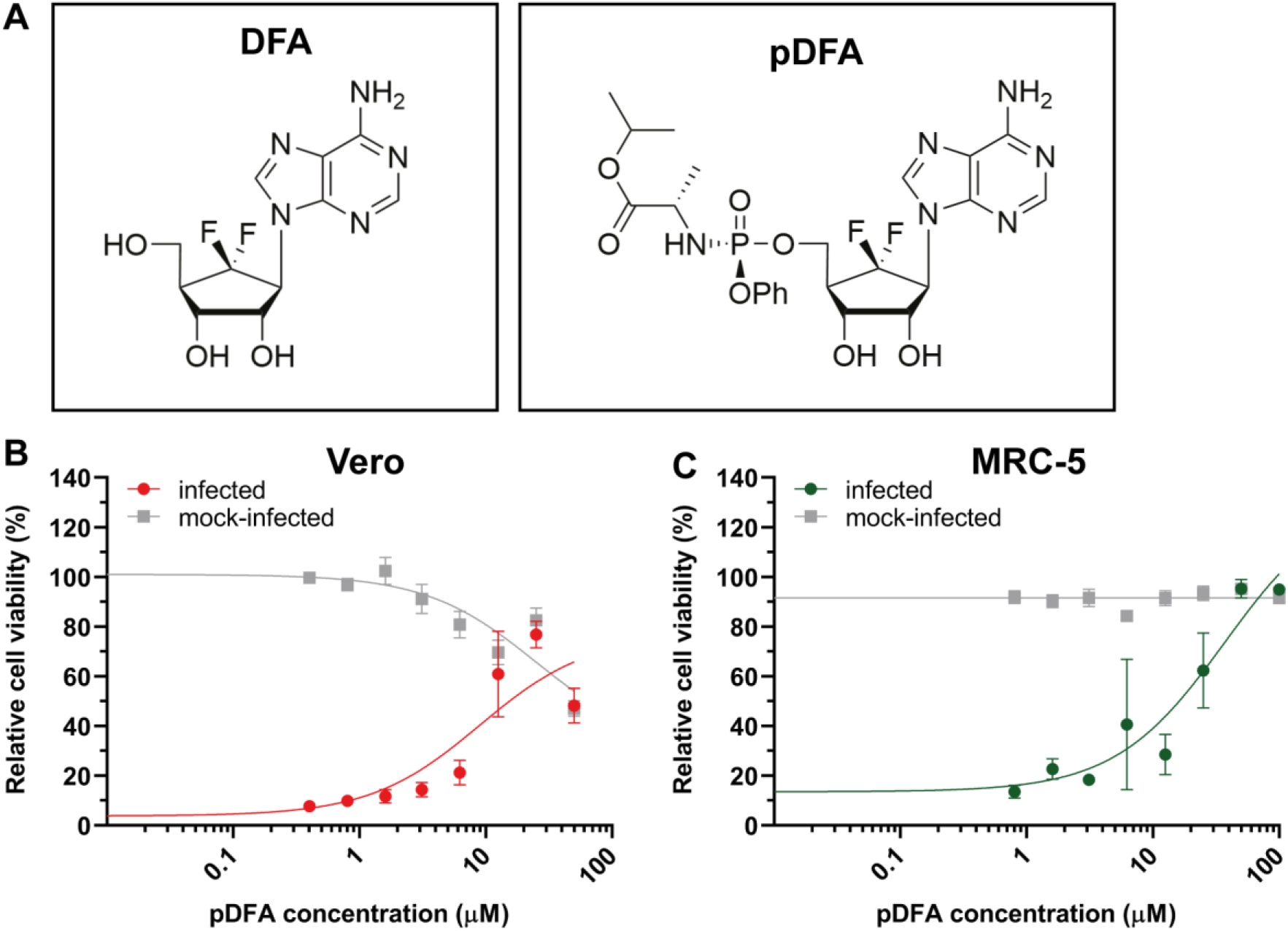
DFA prodrug inhibits MERS-CoV replication. (A) DFA and pDFA schematic structure. pDFA antiviral activity was evaluated by a CPE-reduction assay. (B) Vero or (C) MRC-5 cells were treated with two-fold serial dilution of pDFA and infected with MERS-CoV. After 3 d p.i., cell viability was measured using the CelITiter 96 Aqueous One Solution cell proliferation assay (MTS assay). The graphs show the results of two independent experiments (mean ± sd are shown). A non-linear regression analysis was applied.

### DFA inhibits the early stage of MERS-CoV replication

To characterize the mechanism of action of DFA in more detail, a time-of-addition assay was performed to determine which stage of the viral replication cycle was inhibited by the compound. For this purpose, Vero or MRC-5 cells were infected with MERS-CoV at high MOI (3 PFU/cell) and treated with DFA at different time points pre and post infection at a concentration equaling 4 times the EC_50_. We observed inhibition of replication when the compound was administered before infection and at time points up to 4 h p.i. in Vero (Fig. 3A) and 8 h p.i. in MRC-5 (Fig. 3B). In Vero cells, a 2 log_10_ reduction of progeny virus titers was observed when the compound was administered between 24 h before infection and 1 h p.i. In MRC-5 cells, DFA treatment led to a larger decrease of viral progeny production, >3 log_10_, when treatment was started up to 4 h p.i. Using this DFA dose, no cytotoxicity was detected in either cell line (data not shown). Replication kinetics of MERS-CoV is similar in Vero and MRC-5 cells (48, 49). Thus, the different levels of progeny titer reduction observed between the two cell lines may be explained by variation in uptake or metabolic conversion of the compound (50). In any case, these results demonstrated that DFA inhibits an early stage of MERS-CoV replication.

**Figure 3.**
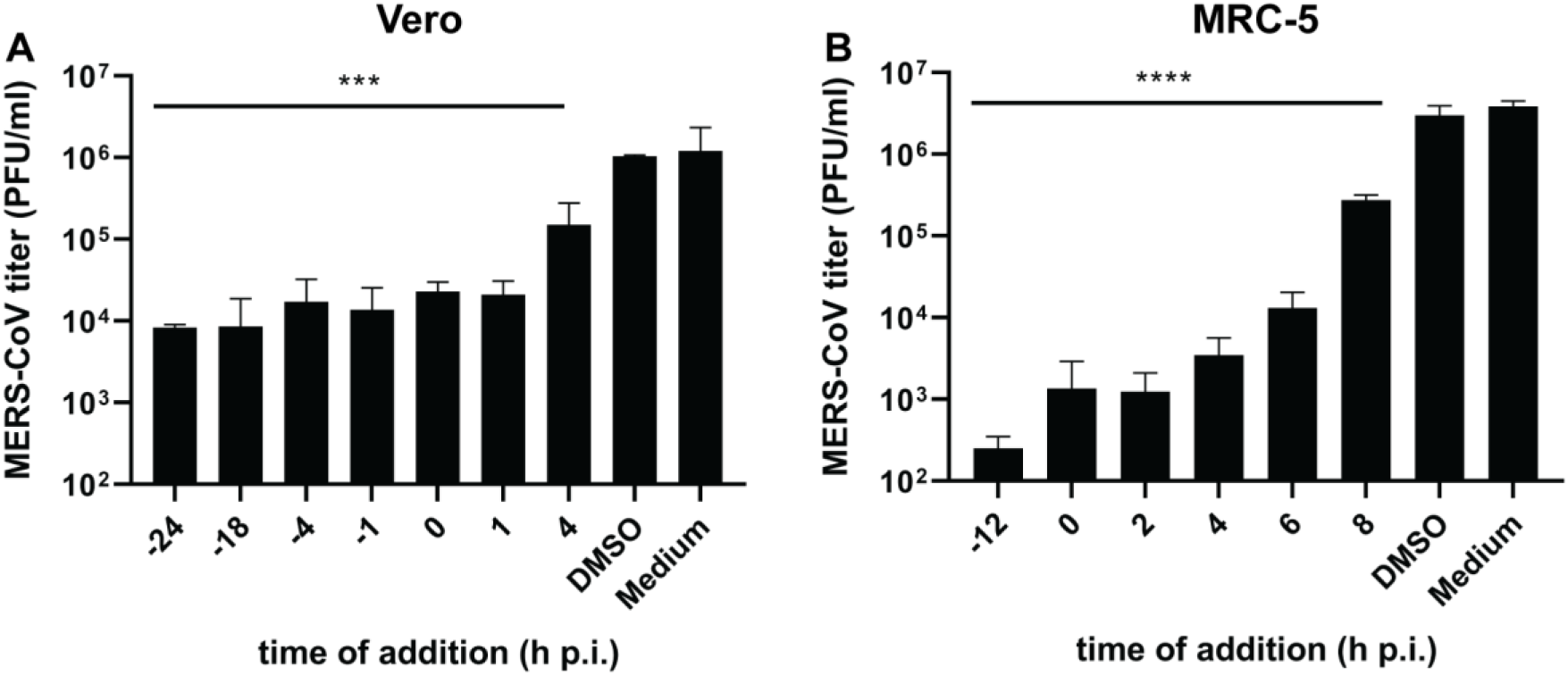
DFA inhibits early steps of MERS-CoV replication. Vero (A) and MRC-5 (B) cells were treated with 0.6 and 12.5 µM, respectively, at the indicated time points pre- and post-infection. Viral progeny in supernatant harvested at 16 h p.i. was determined by plaque assay in Vero cells. The data represent the results from duplicates of 2 independent experiments. Error bars represent standard deviations. Statistical significance was determined by one-way ANOVA.; *, p<0.1; **, p<0.01; ***, p<0.001; ****, p<0.0001.

### Selection of MERS-CoV mutants with 100-fold increased DFA resistance

In order to explore the mode of action of DFA, we selected for compound-resistant MERS-CoV mutants. For this purpose, wild-type MERS-CoV (wtP0) was serially passaged 10 times in MRC-5 cells in the presence of increasing DFA concentrations (from 2.5 µM up to 45 µM). Development of CPE and drug resistance were monitored microscopically, and plaque phenotype and viral progeny production were evaluated by plaque assay after each passage. From passage 8 (P8) onwards, two of the three independently generated lineages showed no increased CPE compared to uninfected, DFA-treated control cells, meaning that these virus populations could not replicate in the presence of DFA concentrations above 35 µM, which were used in these later passages. When P8 virus from lineages 1 or 2 was tested in a CPE-reduction assay, no increased DFA resistance was noticed compared to an untreated wt virus control (data not shown). In contrast, lineage 3 (L3) virus did show clear signs of developing DFA resistance. After 10 passages, infection of cells with L3P10 virus in the presence of 45 µM DFA led to full CPE, which developed equally fast as for the untreated wt control. When L3P10 virus was tested in a dose response assay, only a small (<0.5 log_10_) effect of DFA treatment on viral progeny production was observed in the presence of up to 100 µM of DFA (Fig. 4A). When compared to untreated wt (wtP10) or parental virus (wtP0), L3P10 virus displayed a more than 100-fold increased drug resistance, with an EC_50_ value >100 µM against 0.8 µM for wtP10 and 0.4 µM for wtP0.

**Figure 4.**
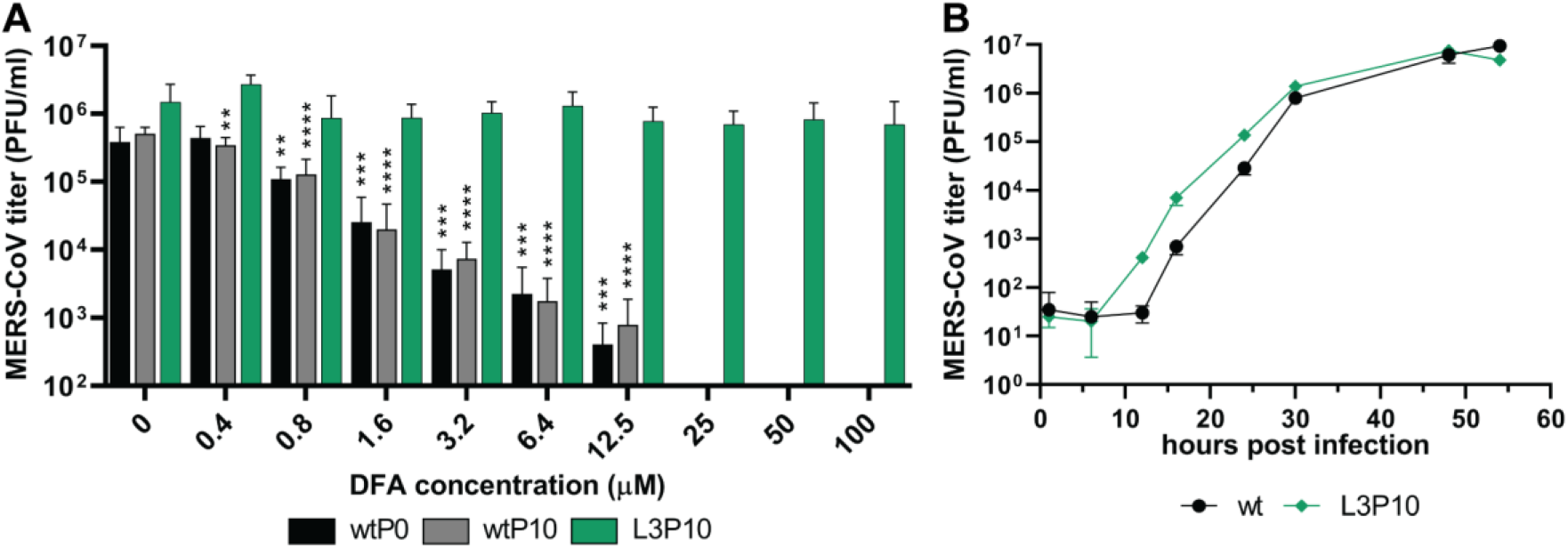
Resistant MERS-CoV mutants selected by passaging in the presence of DFA. Replication in MRC-5 cells of a DFA-resistant virus population (L3P10) in the presence of increasing concentrations of DFA, compared to the parental virus (wtP0) and untreated wt virus (wtP10). Cells were infected with MOI 0.01 and virus progeny in supernatant harvested at 48 h p.i. The data represent the results from four replicates obtained in 2 independent experiments. (B) Characterization of growth kinetics of selected resistant mutant (L3P10). MRC-5 cells were infected with MOI 0.01 and supernatants were harvested at indicated time points from triplicate wells. Viral progeny titers were determined by plaque assay in Vero cells (mean ± sd is presented). Statistical significance was determined by one-way ANOVA. *, p<0.1; **, p<0.01; ***, p<0.001; ****, p<0.0001.

In order to assess if the increased DFA resistance of L3P10 affected its replication kinetics in comparison to the wt control, multi-cycle infections of MRC-5 cells were performed. The two viruses showed similar growth kinetics (Fig. 4B) with peak titers of 6.1×10^6^ PFU/ml (wt) and 7.5 ×10^6^ PFU/ml (L3P10) at 48 h p.i. Taken together, the replication kinetics and strongly increased DFA resistance suggested that, during serial passaging in the presence of DFA, the L3P10 virus population had acquired mutations that account for a strongly increased resistance to the compound.

### Mutations in the L3P10 virus population implicate DFA in the inhibition of viral capping

In order to identify mutations that contribute to DFA resistance, we sequenced the wtP10 and L3P10 virus populations by Illumina next-generation sequencing. Subsequently, sequencing reads were mapped to the reference sequence of MERS-CoV strain EMC/2012 (NC_019843.3; (3)). Sequence variants constituting less than 10% of the total population of viral reads were excluded from further analysis. Compared to the original viral sequence, a total of 14 changes were identified: five synonymous and nine non-synonymous mutations distributed across genes encoding nine different viral proteins. Two of the identified non-synonymous mutations were present both in wtP10 and L3P10, suggesting they were cell culture adaptations acquired during repeated passaging. These mutations, G12033-to-A and C21068-to-U, resulted in D73N and L56F substitutions in nsp7 and nsp16, respectively. Likewise, the five translationally silent mutations in the L3P10 population were considered unlikely to be relevant for its phenotypic profile. Of the remaining seven (Table 1) non-synonymous mutations, one mapped to the accessory protein encoded by ORF5, which is not essential for viral replication in cell culture (51)-(52), and one to the Spike protein (53), which also is an unlikely target for inhibition by nucleoside analogues. This left five L3P10-specific mutations leading to amino acid substitutions in the viral replicase subunits nsp1, 3, 12, and 13 that may be associated with DFA resistance. As shown in Table 1, all of these were present in only part of the viral population (in 37% to 55% of the total reads), suggesting a complex pattern of virus evolution with DFA resistance possibly relying on (different) combinations of mutations (see Discussion). The short NGS read-length (150 nucleotides) did not allow us to determine which mutations were combined in the same genome. Since further optimization of this compound class is needed to improve its selectivity index (Table 2), we did not perform follow-up experiments to elucidate its mode of action at this stage.

**Table 1.** Summary of non-synonymous mutations specifically identified in MERS-CoV L3P10 by NGS

| Coding region | nt change | aa change | Domain | Presence in L3P10 (NGS) |
| --- | --- | --- | --- | --- |
| **nsp1** | G795C | D172H | CTD | 55% |
| **nsp3** | C6777T | R1314C | PLnc | 37% |
| **nsp12** | A14061T | Y218F | NiRAN | 48% |
| **nsp13** | G16272A | R22K | ZBD | 49% |
|  | G16689A | R161H | 1B | 40% |
| **spike** | T24085_24086insACTCAACAGGTG | P876_V877ins<br>TQQV |  | 37% |
| **ORF5** | G26927_26928insT | Not in frame |  | 48% |

nt, nucleotide; aa, amino acid; CTD, C-terminal domain; PLnc, papain-like noncanonical domain; NiRAN, Nidovirus RdRp associated nucleotidyl transferase domain; ZBD, Zinc binding domain; 1B, 1B domain of helicase; \* also present in wtP10 control

**Table 2.** The antiviral effect of DFA on the replication of different betacoronaviruses.

| **Virus** | **Cell line** | **EC<sub>50</sub> (μM)** | **CC<sub>50</sub> (μM)** | **SI** |
| --- | --- | --- | --- | --- |
| **MERS-CoV** | Vero | 0.2 | >3.2 | >16 |
|  | HuH7 | 0.6 | >50 | >80 |
|  | MRC-5 | 0.8 | >50 | >60 |
|  | Calu-3 | ~2 | >25 | >12 |
| **SARS-CoV** | Calu-3 | ~5 | >25 | >5 |
| **SARS-CoV-2** | Calu-3 | ~2 | >25 | >12 |

EC<sub>50</sub>s values were calculated based on results obtained in dose response assay, while CC<sub>50</sub>s values were determined in a cell viability assay as described in materials and methods. SI, selectivity index was calculated by comparing CC<sub>50</sub> with EC<sub>50</sub> values.

### Evaluation of DFA potential as a pan-coronaviral inhibitor

To explore the potential of DFA as a broad-spectrum antiviral, Calu-3 cells, human lung cells that supports MERS-CoV, SARS-CoV and SARS-CoV-2 replication (54, 55), were treated with increasing concentrations of DFA and infected with each of these viruses in a dose response assay. By using the same cell line for all three CoVs, differences in DFA up-take or metabolic conversion to its triphosphate form were eliminated. The results showed a dose-dependent decrease in the production of viral progeny for MERS-CoV (Fig. 5A) and SARS-CoV-2 (Fig. 5C) that followed the cytotoxicity of the compound. At a DFA concentration of 3.2 µM, only a small reduction of MERS-CoV and SARS-CoV-2 progeny was observed, 0.5 to 1log_10_. Surprisingly, the antiviral activity of DFA against MERS-CoV in Calu-3 cells was severely reduced when compared to results obtained in other cell lines, including another human lung cell line MRC-5 (Fig. 1D-F). In the case of SARS-CoV infection, a minor inhibitory effect was observed at concentrations that appeared to be somewhat cytotoxic (Fig. 5B and 5D), contrary to what was demonstrated in Vero E6 cells ((38) and Table 2). Unfortunately, in Calu-3 cells cytotoxicity was detected at low compound concentrations (>6.2 µM) and the inhibitory effects observed could thus be associated with an overall decrease in relative cell viability. This indicates that the design of improved DFA derivatives is needed to decrease cytotoxicity and improve inhibitory potency.

**Figure 5.**
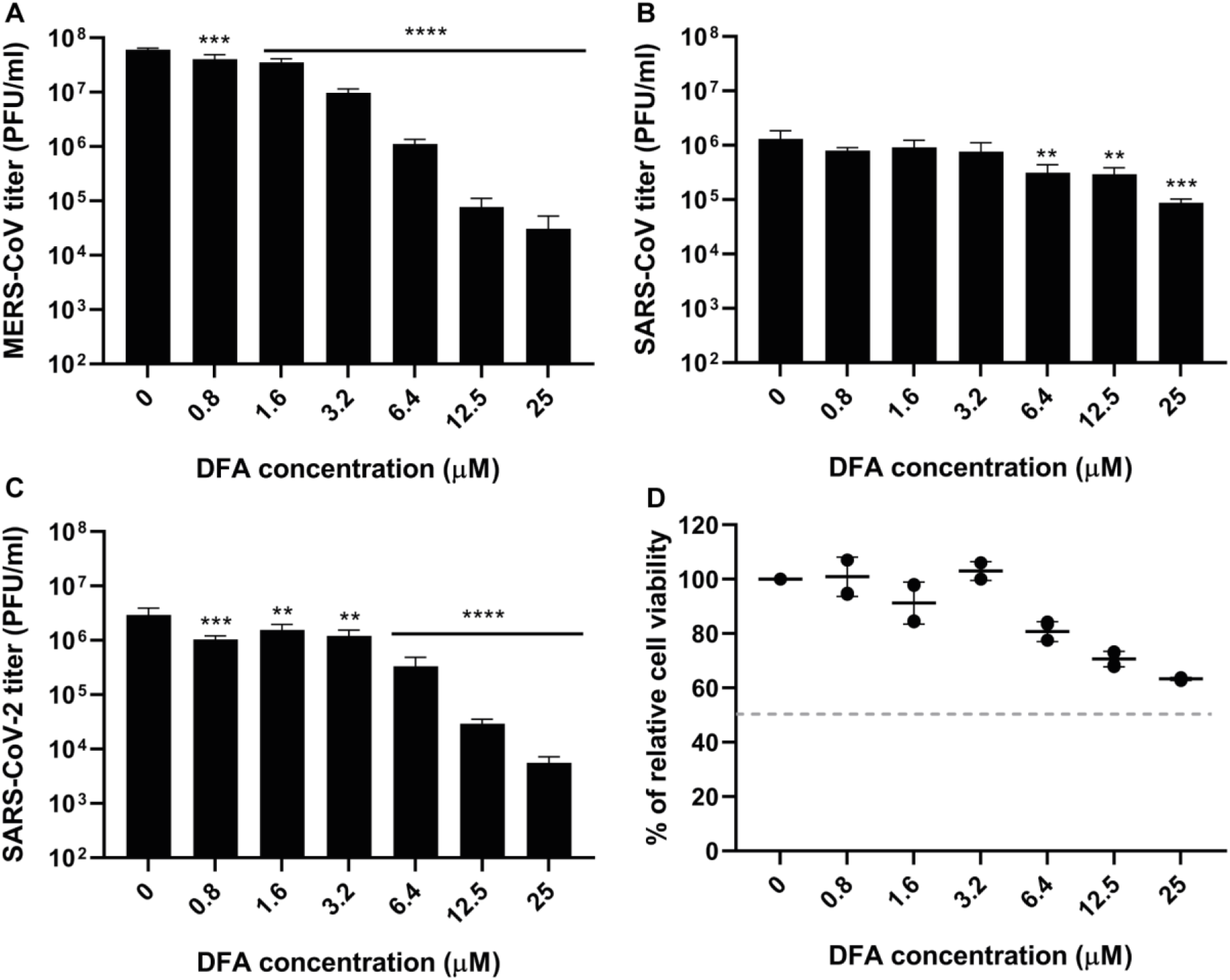
DFA antiviral activity is reduced in Calu-3 cells. Calu-3 cells were infected with MERS-CoV (A), SARS-CoV (B) and SARS-CoV-2 (C) in the presence of various concentrations of DFA. An MOI of 1 was used, based on titrations of virus stocks on Vero cells. Progeny virus titers in supernatants harvested at 24 h p.i. were determined by plaque assay in Vero cells. (D) Cytotoxicity of DFA was measured in mock-infected cells, and was determined at 24 h p.i. in a CPE-reduction assay by use of the CelITiter 96 Aqueous One Solution cell proliferation assay (MTS assay). The data represent triplicates of 2 independent experiments and error bars show standard deviations. Statistical significance was determined by one-way ANOVA. No*, no significance; *, p<0.1; **, p<0.01; ***, p<0.001; ****, p<0.0001.

## Discussion

This study describes that treatment with low-micromolar DFA concentrations exhibits a strong antiviral effect on MERS-CoV replication in cell culture-based infection models (Fig. 1). Time-of-addition assays indicated that DFA reduced MERS-CoV progeny production when cells were treated prior to, at the time of, or within 4 h after infection (Fig. 3), suggesting that DFA interferes with the early stage of replication. Propagation of MERS-CoV in the presence of DFA led to the selection of a virus population with strongly enhanced resistance to this compound (Fig. 4). Subsequent sequence analysis revealed a potentially complex pattern of resistance evolution, exhibiting multiple mutations that are present in only part of the virus population, including several that map to enzymes involved in viral RNA synthesis and mRNA capping (Table 1).

DFA was originally designed to target the host SAH hydrolase directly and was demonstrated to inhibit this enzyme *in vitro* with an IC_50_ (50% inhibitory concentration) of 1.06 µM (38). The compound is a carbocyclic adenosine analogue based on the parental inhibitor aristeromycin (56, 57), which was further modified by incorporation of a difluorine group at the 6’ (top) position of its sugar ring ((38) and Fig. 2A). This modification improved the binding affinity of the compound for human SAH hydrolase and, consequently, the inhibition of its enzymatic activity. Previous studies demonstrated that treatment of cells with high-affinity SAH hydrolase inhibitors, such as neplanocin A and aristeromycin, increases the intracellular SAH concentration, preventing the metabolic conversion of SAH to adenosine and L-homocysteine (reviewed in (58)). Therefore, SAH hydrolase inhibitors reduce or deplete the intracellular pools of homocysteine and adenosine, the latter being produced exclusively by SAH hydrolysis. As the SAM methyl donor is formed via homocysteine trans-sulfuration or the adenosine kinase pathway, SAH hydrolase regulates the intracellular SAM levels and consequently the cell’s SAM-dependent methylation reactions. Moreover, SAH accumulation can also reduce the activity of SAM-dependent methyltransferases by feed-back inhibition, as SAH can bind to their active site with higher affinity than SAM itself (36).

A correlation between the antiviral effect of adenosine analogues and their ability to interfere with viral capping has been demonstrated in previous studies with chikungunya virus, dengue virus, West Nile virus and vaccinia virus (59-62). Both CoV methyltransferases use SAM as a methyl donor for their enzymatic activity (63-65). Thus, SAH hydrolase inhibition and reduced SAM concentrations may impact, if not block, their activity. Previous studies with 5’-β-fluoroadenosine and derivatives of aristeromycin demonstrated that inhibition of SAH hydrolase affects viral replication by reducing RNA methylation (reviewed in (58)). Biochemical analysis of the MERS-CoV nsp16/nsp10 complex showed the capacity of nsp16 to bind SAH with greater affinity than SAM (66), whereas superimposition of the SARS-CoV nsp16/nsp10 in complex with SAH demonstrated that the same binding site is used by both substrates (67). In addition, increased SAH concentrations reduced the 2’-O-methylation of N7-methylated substrates (65, 66).

In light of the above, direct or indirect inhibition of the CoV capping pathway is a possible mode of action (MoA) of DFA, although the genotypic profile of the L3P10 virus population (Table 1) suggests that the compound may inhibit CoV replication using multiple mechanisms. As DFA is a nucleoside analog, mutations in viral enzymes involved in RNA synthesis and capping may contribute to the observed DFA-resistance of the L3P10 virus population. Consequently, mutations Y218F in the nsp12-NiRAN domain and R22K and R161H in the nsp13 ZBD-helicase subunit attract more attention than nsp1-D172H and nsp3-R1314C. Recently, the nsp12 NiRAN domain was proposed to function as the capping GTase (31), while also nsp13 has been implicated in the CoV capping pathway (see Introduction; reviewed in (30)). As the identified mutations have not been characterized in structural or biochemical studies, one can only speculate about their potential role in viral replication and DFA resistance

Further phenotypic and mechanistic studies will be needed to better understand the mode of action of DFA. Additionally, cloning of L3P10 viruses by plaque picking could help to define the combination(s) of mutations that are the basis for DFA resistance, by evaluating their frequency of occurrence and associated replication and plaque phenotype.

As a nucleoside analogue, DFA was also considered to be a potential RdRp inhibitor. This would require uptake by the cell’s nucleoside transporters, and subsequent phosphorylation into a triphosphorylated product that could be incorporated into the RNA chain during viral RNA synthesis (reviewed on (47)). In order to improve absorption of the compound by the cells and metabolism into its active form, a prodrug of DFA was synthesized and its antiviral activity was evaluated. In theory, the monophosphoramidate mask would promote the second phosphorylation to occur once the compound enters the cytoplasm by circumventing the rate-limiting step of the first phosphorylation. However, when compared to DFA, the EC_50_ of the prodrug was more than 10 times higher (Fig. 2), in contrast to results obtained with prodrugs of other nucleoside analogues (46, 68). In previous work, structure-activity studies and tests of several purine and pyrimidine analogues of DFA suggested that DFA is most likely not targeting the RdRp (38, 61, 69). This notion is also supported by the fact that the genotypic profile obtained for L3P10 did not reveal mutations in the RdRp domain of nsp12.

In this study, we demonstrate that DFA can inhibit the replication of MERS-CoV, but that the design and development of DFA-based derivatives will be required to reduce cytotoxic side effects. Combining our results in this study with our previous report (38), showing that DFA can inhibit chikungunya and Zika virus, DFA appears to be an interesting compound for further development as a broad-spectrum antiviral agent.

## Materials and Methods

### Cell culture and viruses

Vero cells were a kind gift from the Department of Viroscience, Erasmus Medical Center, Rotterdam, the Netherlands, and Huh7 cells were provided by Dr. Ralf Bartenschlager, Heidelberg University, Germany. Vero, Vero E6, Huh7, MRC-5 and Calu-3 were cultured as described before (48, 49, 70-72). All cells were incubated at 37°C with 5% CO_2_. Infections were carried out in Eagle’s minimum essential medium (EMEM; Lonza) containing 25 mM HEPES (Lonza), 2% fetal calf serum (FCS; Bodinco), 100 units/ml penicillin (Lonza), 100 units/ml streptomycin (Lonza) and 2 mM L-glutamine (PAA) (abbreviated from now on as EMEM-2%FCS). MERS-CoV (strain EMC/2012; (3, 4)), SARS-CoV (Frankfurt-1 strain,(73)) and SARS-CoV-2/Leiden-0002 (GenBank accession nr. MT510999; (72)) were used for infections with wild type virus. CoV infections were performed inside biosafety cabinets in a certified biosafety level 3 (BSL3) facilities at Leiden University Medical Center.

### Compounds

6′,6′-Difluoro-aristeromycin (DFA) and its adenine phosphoramidate pro-drug (pDFA) were designed and synthesized, designated as 2c and 3a, respectively, as described in a previous report (38). Different batches of powder were dissolved in DMSO to a final concentration of 20 mM and single use aliquots were stored at 4°C. Remdesivir (RDV; HY-104077) was purchased from MedChemexpress and chloroquine (C6628) from Sigma. Both compounds were dissolved in adequate solvents (DMSO or PBS, respectively) and single use aliquots were stored at −20°C.

### Cytopathic effect (CPE) reduction assay

Cells were seeded in 96-well flat bottom plates in 100 µl at a density of 10000 cells/well of Huh7, 15000 cells/well of MRC-5 or 20000 cells/well of Vero cells. After overnight culture at 37°C, cells were pre-incubated for 30 min with 50 µl of two-fold serial dilutions of compounds prepared in EMEM-2%FCS. Subsequently, half of the wells were infected with MERS-CoV at low MOI in a total volume of 150 µl of medium with increasing concentrations between 0.05 to 100 uM of compound, to evaluate the inhibitory effect of compound. The other half of the wells were “mock”-infected with medium to monitor the (potential) cytotoxicity of the compound. Plates were incubated for three days (or as mentioned) at 37°C, after which cell viability was measured using the colorimetric CellTiter 96® Aqueous Non-Radioactive Cell Proliferation kit (Promega). The absorption at 495 nm was measured using a monochromatic filter in a multimode plate reader (Envision; Perkin Elmer). Data were normalized to the “mock”-infected control, after which EC_50_ and CC_50_ values were calculated using non-linear regression with Graph-Pad Prism V8.0. Each experiment was performed at least in quadruplicate and repeated at least twice.

### Dose response assay

To evaluate the effect of compound treatment on viral progeny titers, confluent monolayers of Vero, Huh7 or MRC-5 were seeded in 24-well plates. Cells were incubated for 30 min at 37°C with solvent or a range of DFA concentrations (from 0.1 to 100 uM). Then, cells were infected with MERS-CoV at an MOI of 0.01 for 1 h. After infection, cells were washed three times with PBS and 1 ml of medium with compound at corresponding concentration was added. Supernatants were collected at 48 h p.i. and viral progeny titers were determined by plaque assay in Vero cells as described before (74).

In 96-well clusters, Calu-3 cells were seeded at a density of 3 × 10^4^ cells per well in 100 µl culture medium. Two days later, cells were pre-incubated for 30 min with 2-fold serial dilutions of compound, starting at a concentration of 25 µM. Subsequently, cells were infected with MERS-CoV, SARS-CoV or SARS-CoV-2 (MOI of 1 based on titer determined on Vero cells) in the presence of compound for 1 h. Next, cells were washed three times with PBS and 100 µl of compound solution in EMEM-2%FCS was added. Supernatants were collected at 24 h p.i. and progeny virus titers were determined by plaque assay.

### Time of addition assay

Confluent monolayers of MRC-5 or Vero cells were seeded in 12-well plates in 1 ml/well of the appropriate medium (see above), and were grown overnight at 37°C. Treatment of cells (before, during or after infection) was performed using 0.6 µM (for Vero) and 12.5 µM (for MRC-5) of compound solution freshly prepared in EMEM-2%FCS medium. Cells were infected with MERS-CoV inoculum (MOI of 5) for 1h and washed three times with PBS. Subsequently, EMEM-2%FCS medium was added to the cells and supplemented with compound solution in 2-h intervals to a final concentration as mentioned above. Supernatants were collected 16 h p.i. and viral titers were determined by plaque assay.

### Resistance culturing and next-generation sequencing (NGS)

Recombinant wt MERS-CoV strain EMC/2012(rMERS-CoV) was passaged in triplicate in presence of increasing concentrations of DFA ranging from 3.2 µM to 45 µM. Infections were performed at an MOI of 0.05 in every passage in MRC-5 monolayers. In parallel, rMERS-CoV wt was passaged in the same conditions in the absence of compound, to identify possible mutations associated with cell culture adaptation. Additionally, a “mock”-infected well treated with the same concentration of compound in each passage was evaluated for cytotoxicity by light microcopy. Supernatants were harvested when 80% to full CPE was observed (usually at 3 d p.i.). Three lineages were generated by serial passaging, but only lineage 3 was used for next-generation sequencing. To this end, RNA was isolated from 200 µl of virus-containing cell culture supernatants using TriPure isolation reagent (Roche Applied Science) and purified according to manufacturer’s instructions. The RNA concentration was measured using a Qubit fluorometer and RNA High Sensitivity kit (Thermo Fisher Scientific). NGS sample preparation and analysis were performed as described previously (75). After filtration and trimming of data, the remaining reads were mapped to the MERS-CoV GenBank reference sequence (NC_019843;(3, 4)). Raw NGS data sets for wtP10 and L3P10 samples analysed in this study were deposited in NCBI Bioproject and are available under the following link: http://www.ncbi.nlm.nih.gov/bioproject/730836. Only MERS-CoV-specific reads were included in these data files.

## Acknowledgments

N.S.O. was supported by the Marie Skłodowska-Curie ETN European Training Network ‘ANTIVIRALS’ (EU Grant Agreement No. 642434). We thank Adriaan de Wilde and Martijn van Hemert for their support and for helpful discussions. We thank Igor Sidorov for his technical support in the analysis of NGS data.

## Notes

### Competing Interest Statement

The authors have declared no competing interest.

### Summary of Updates

- sequencing data. - shortened the theoretical part. - updated Figure 3 and Table 1.

